# netCRS: Network-based comorbidity risk score for prediction of myocardial infarction using biobank-scaled PheWAS data

**DOI:** 10.1101/2021.10.12.464134

**Authors:** Yonghyun Nam, Sang-Hyuk Jung, Anurag Verma, Vivek Sriram, Hong-Hee Won, Jae-Seung Yun, Regeneron Genetics Center, Dokyoon Kim

## Abstract

The polygenic risk score (PRS) can help to identify individuals’ genetic susceptibility for various diseases by combining patient genetic profiles and identified single-nucleotide polymorphisms (SNPs) from genome-wide association studies. Although multiple diseases will usually afflict patients at once or in succession, conventional PRSs fail to consider genetic relationships across multiple diseases. Even multi-trait PRSs, which take into account genetic effects for more than one disease at a time, fail to consider a sufficient number of phenotypes to accurately reflect the state of disease comorbidity in a patient, or are biased in terms of the traits that are selected. Thus, we developed novel network-based comorbidity risk scores to quantify associations among multiple phenotypes from phenome-wide association studies (PheWAS). We first constructed a disease-SNP heterogeneous multi-layered network (DS-Net), which consists of a disease network (disease-layer) and SNP network (SNP-layer). The disease-layer describes the population-level interactome from PheWAS data. The SNP-layer was constructed according to linkage disequilibrium. Both layers were attached to transform the information from a population-level interactome to individual-level inferences. Then, graph-based semi-supervised learning was applied to predict possible comorbidity scores on disease-layer for each subject. The SNP-layer serves as receiving individual genotyping data in the scoring process, and the disease-layer serves as the propagated output for an individual’s multiple disease comorbidity scores. The possible comorbidity scores were combined by logistic regression, and it is denoted as netCRS. The DS-Net was constructed from UK Biobank PheWAS data, and the individual genetic profiles were collected from the Penn Medicine Biobank. As a proof-of-concept study, myocardial infarction (MI) was selected to compare netCRS with the PRS with pruning and thresholding (PRS-PT). The combined model (netCRS + PRS-PT + covariates) achieved an AUC improvement of 6.26% compared to the (PRS-PT + covariates) model. In terms of risk stratification, the combined model was able to capture the risk of MI up to approximately eight-fold higher than that of the low-risk group. The netCRS and PRS-PT complement each other in predicting high-risk groups of patients with MI. We expect that using these risk prediction models will allow for the development of prevention strategies and reduction of MI morbidity and mortality.

## 1. Introduction

The prediction of an individual’s disease risk is an essential part of precision medicine and will be required to improve public healthcare and understand risk of developing a disease across different populations. One of the most popular methods of disease risk prediction is the polygenic risk score (PRS), which estimates a patient’s genetic risk for a chosen trait or disease by combining individual genetic profiles with many single-nucleotide polymorphisms (SNPs) identified through genome-wide association studies (GWAS).^1,2^ Many studies have calculated PRSs for various common diseases, including cardiovascular disease, hypertension, and neurological disorders, and they suggest that the PRS might be a helpful tool for identifying and categorizing high-genetic risk individuals for those diseases.^3-6^ Nevertheless, a major weakness of the conventional PRS is its focus on a single trait for the estimation of genetic risk scores – when predicting the risk scores of an index disease of interest, PRS is calculated according solely to the relevant phenotype. In most cases, however, multiple diseases will usually afflict a patient at once or in succession. These disease complications and comorbidities, referring to the presence of one or more additional medical conditions given a primary disease, suggest that effective disease prediction will require us to consider multiple phenotypes concurrently.^7^ In order to estimate the disease risk considering the associations among multiple diseases, several studies had attempted to perform the association analysis for PRSs with multiple diseases through subsequent analysis^8,9^ or to combine PRSs for multiple traits.^10^ In these previous studies, a key step involves the determination of which diseases related to the index disease are selected for estimation of the combined risk score. However, these methods are limited as selection bias is introduced when knowledge reveled in clinical practice is used to identify diseases highly related to the target phenotype. Even multi-trait PRSs, which take into account genetic effects for more than one disease at a time, fail to consider a sufficient number of phenotypes to accurately reflect the state of disease comorbidity in a patient, or are biased in terms of the traits that are selected.

One effective way to comprehensively explore the genetic associations among multiple diseases is to consider a network representation, such as the disease-disease network (DDN). Given a set of diseases, the DDN represents diseases as nodes, and disease-disease associations as edges. DDNs can explore potential comorbidity relationships among phenotypes based on shared genetic components. Different genetic components will yield different types of networks, such as gene^11^, protein^12^, pathway^13^, and SNP-based DDN.^14^ In this study, the SNP-based DDN is used to incorporate the conventional PRS approach, where edges represent the number of shared SNPs between diseases according to results from a phenome-wide association study (PheWAS). The SNP-based DDN using PheWAS results is depicted in the center panel of Figure 1. Considering D2 as an index disease of interest (marked in red), we can see that it is directly connected with four diseases (D1, D3, D4, and D6). Three diseases (D5, D7, and D8) share no edges with D2. Directly connected diseases share at least one common SNP with D2. Indirectly connected diseases share no genetic associations with D2, but they are connected through the other nodes – for instance, D2 and D7 are connected in through the sequence of diseases with D2∼D6∼D7. Overall population-level relationships between diseases can be observed through the underlying structure of the DDN, regardless of whether or not a pair of diseases share genetic components. In developing risk prediction models which consider the relationships across a multitude of diseases, a DDN can provide intuitive, unbiased evidence about the selection of related diseases as well as the strength of associations between diseases. However, although the population-level interactome between phenotypes can be observed through a DDN, it is not easy to apply these disease-disease associations in a patient-specific manner. Indeed, it is difficult to obtain information pertinent to the individual because the nodes and edges in DDN are aggregated and summarized from PheWAS data.

**Figure 1.**
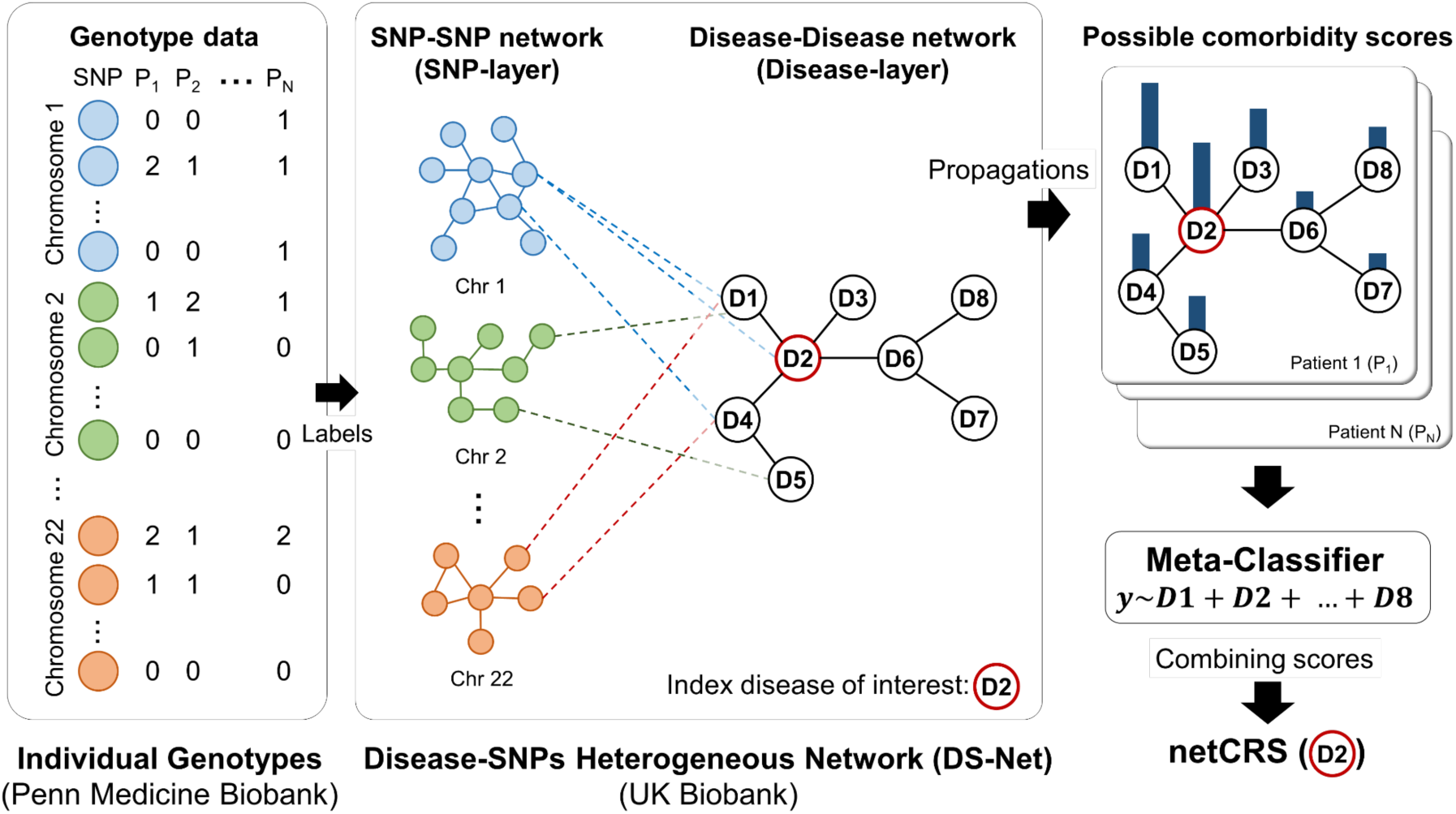
Overall framework of network-based comorbidity risk scoring algorithms (netCRS): **Left)** individual genotype data collected from Penn Medicine BioBank. **Middle)** schematic description of disease-SNP heterogeneous multi-layered network (DS-Net). SNP-layer constructed by linkage-disequilibrium and disease-layer constructed using UK biobank PheWAS summary data. **Right)** Upper right represents possible comorbidity scores of each disease for individual. The possible comorbidity scores are combined by logistic regression, and the combined scores, netCRS, are generated by each patient

To circumvent this challenge, we propose a novel framework of network-based individual comorbidity risk scores (netCRS) to predict individual-level disease comorbidity risk through population-level interactome networks. The goals of netCRS are as follows: (a) To improve the prediction ability of PRS, we present a novel risk score that estimates multiple disease comorbidities according to their shared genetic components. The netCRS estimates the combined comorbidity scores for multiple phenotypes in the SNP-based DDN when provided with an individual genetic profile. In PRS, marginal effect size estimates of SNPs obtained from a GWAS are used as weights for weighted sum scores of risk alleles carried by an individual for a single trait. On the other hand, in netCRS, disease-specific effect size estimates of SNPs from PheWAS are used as edge weights of the network for multiple traits. (b) To obtain individual-level inference from population-level interactome, we construct a novel disease-SNP heterogeneous multi-layered network using EHR-linked biobank-scale PheWAS summary statistics. Using this multi-layered network, we introduce a scoring method to infer individual information from population-level networks through layer-wise label propagation.

Figure 1 describes the overall conceptual framework of netCRS. The center panel depicts a disease-SNP heterogeneous multi-layered network (denoted as DS-Net). The DS-Net is a multi-layered graph, consisting of a SNP-SNP correlation network (SNP-layer), disease-disease network (disease-layer) and SNP-disease associations (coupling graphs). Briefly, the SNP-layer (colored solid circles/lines) is constructed according to a linkage disequilibrium matrix, and the disease-layer (colorless solid circles/lines) is constructed according to the shared genetic components between phenotypes. The coupling graphs for inter-layers (colored dashed lines) between the SNP- and disease-layer are derived using disease-SNP associations obtained from PheWAS summary statistics. Given the DS-Net and index disease of interest, we first predict individual comorbidity scores using graph-based semi-supervised learning (SSL). Graph-based SSL predicts scores on the disease-layer by propagating label information when the individual genetic profile is labeled on the SNP layer. In the left panel of Figure 1, individual genotype data is used to provide query or seed label information to the SNP-layer for the scoring algorithm. Each patient’s genetic data are initially labeled on the SNP-layer, and then the label information is propagated through the multi-layered network. Predicted risk scores are obtained for each disease node (blue bar). Each bar depicts a possible comorbidity score for each disease that an individual patient can have. The predicted comorbidity scores are subsequently aggregated into combined comorbidity scores using a meta-classifier (the right panel of Figure 1). Here, we use logistic regression for our meta-learner, and the combined comorbidity score is denoted as netCRS(), where the parentheses specify the index disease of interest. More details of the proposed methods are explained in the following sections.

## 2. netCRS: Network-based individual Comorbidity Risk Scoring

### 2.1. Disease-SNP Heterogeneous Network using UK Biobank summary statistics

We constructed the reference network using UK BioBank (UKBB) PheWAS summary statistics. The DS-Net is a multi-layered weighted graph, ***G*** = (***V, W, S***), where ***V*** represents the set of nodes, ***W*** represents the set of edges, and ***S*** represents the set of layers. The multi-layered network ***G*** is decomposed into two distinct single graphs with corresponding layers *S* = {*S*_Disease_, *S*_SNP_}. The similarity matrix ***W*** for multi-layered network can be expressed in block-wise matrix as follows:

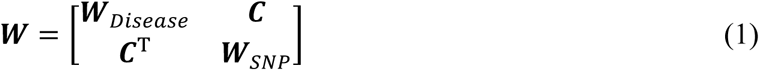

The block diagonal matrix (***W***_Disease_ and ***W***_SNP_) represents a similarity matrix for each single network of the disease-layer and SNP-layer respectively, and the block-off diagonal matrix ***C*** represents the coupling graphs for the connections between inter-layers.

#### 2.1.1. Disease-Layer (Disease-Disease network)

The disease-layer ***G***_Disease_ = (***V***_Disease_, ***W***_Disease_) is a sub-network of the DS-Net ***G***, where the nodes ***V***_Disease_ denotes the set of diseases, and ***W***_Disease_ denotes the similarity between the sequences of SNPs that pairs of diseases commonly share. The disease-layer is constructed according to shared genetic components, with the hypothesis that two different phenotypes are associated if they share significant SNPs from the PheWAS summary results. Given *m* diseases and *k* SNPs, we first generate *m* disease-SNP association vectors from each PheWAS result. Each disease vector ***v*** is represented as a *k*-dimensional SNP vector with binary attributes, each of which stands for statistically significant (‘1’) or not (‘0’) for the association with a specific SNP that has passed the *p*-value thresholds in the PheWAS results.^14^ Then, similarity between pairs of diseases is measured by cosine similarity *w*_*ij*_ for two diseases *v*_*i*_ and *v*_*j*_.

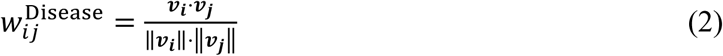

#### 2.1.2. SNP-layers (SNP-SNP correlation network)

SNP-layer ***G***_SNP_ = (***V***_SNP_, ***W***_SNP_) is a sub-network of the disease-SNP heterogeneous network ***G*** when ***S*** = {S_SNP_}. The node ***V***_SNP_ denotes the representative SNPs after genetic pre-processing, and ***W***_SNP_ denotes the pairwise genetic correlations between distinct SNPs. We generate the pairwise linkage-disequilibrium (LD) matrices of genotype correlation between nearby SNPs using quality-controlled genotyped data of UKBB samples. The *r*^2^ between pairs of SNPs is obtained using PLINK 1.90 with LD calculation (--r2, --ld-window 10 SNPs, --ld-window-kb 1000kb, and --ld-window-r2 0.0). The similarity matrix ***W***_SNP_ is composed of correlation values ranging from 0 to 1.

#### 2.1.3. Coupling graphs (SNP-Disease associations)

The coupling graphs ***C*** = {*c*_*ik*_| *i ∈* ***V***_*Disease*_ *k ∈* _*SNP*_} imply connections between diseases and SNPs across different layers of the network. Coupling edges are derived from the disease-SNP association vectors (described in section 2.1.1). Edge weights take value of z-scores, equivalent to the beta-coefficients (***β***_*ik*_) divided by standard errors (SE_*ik*_) from the significant association between phenotype *i* and SNP *k* from PheWAS results. These weights are rescaled to lie within a range of 0 to 1.

Combining the disease-layer, SNP-layer, and coupling graphs yields the proposed DS-Net. The constructed network can provide insights into the population-level interactome between diseases and SNPs.

### 2.2. Individual comorbidity risk scoring algorithms

Given an index disease of interest, we can predict individuals’ disease comorbidity risk scores using the DS-Net. Since the network describes a biobank-scale population-level interactome, we take individual genetic information from another biobank to calculate risk scores for individual patients. In this analysis, the summary-level data from UKBB were used for the network construction, and the individual-level genetic data were collected from the Penn Medicine BioBank (PMBB). More details are explained in the Section 3.

Let us define disease comorbidity risk scoring ***f***: ***V*** →ℝ as a function that quantifies the degree of commitment of the diseases associated with SNPs on the network. To implement this scoring function in a DS-Net, we employ graph-based SSL with transductive learning settings.^15^ As shown in Figure 1, individual genotypes are used for initial label information in the DS-Net. We set the genotype CC (homozygous non-reference) as 0, genotype CT (heterozygous) as 0.5, and genotype TT (homozygous reference) as 1 for initial labels of label propagation. Once the labels for the SNP-layer are provided, graph-based SSL propagates the label information through all edges in the heterogeneous multi-layered network simultaneously. Since we are interested only in the comorbidity risk of multiple diseases, the propagated disease scores ***f***_Disease_ on the disease-layer ***V***_Disease_ are used as the predicted comorbidity feature vectors. To aggregate these scores, we employ logistic regression as the meta-classifier.

The following section describes the formulation of the proposed network-based comorbidity scoring algorithm. Assume that we have genotype data for *m* individuals and that we know the diagnosis outcomes of the index disease. Then, *i*-th patient’s genotype data ***m***_*i*_ has *k*-dimensional SNP vectors with values of 0, 0.5, and 1 as described above. The outcomes of the index disease for all patients ***z*** is an *m*-dimensional vector with value ‘1’ if the patient has been diagnosed with the index disease or ‘0’ otherwise. To apply the individual data to graph-based SSL, we set the initial label set of vector ***y*** and predicted scores ***f***. The initialization and learning process is performed iteratively patient-by-patient. Let ***y*** = (*y*_1_, …, *y*_*n*_, *y*_*n*+1_, …, *y*_*n*+*k*_)^T^= (***y***_Disease_, ***y***_SNP_)^T^ denote the set of initial labels and ***f*** = (*f*_1_, …, *f*_*n*_ *f*_*n*+1_, …, *f*_*n*+*k*_)^.T^ = (***f***_Disease_, ***f***_SNP_)^.T^denote the set of predicted scores, where *n* is the total number of diseases and *k* is the total number of SNPs in the network. In the problem setting of disease comorbidity scores, we set the ***y***_Disease_ to the zero vector and ***y***_SNP_ to ***m***_*i*_. The label information is propagated to all connected nodes along with edges in ***W***_SNP_, ***C***, and ***W***_Disease_ on graph ***G***. Graph-based SSL provides the real-valued scores ***f*** with two assumptions: (a) smoothness function (predicted scores *f*_*i*_ and *f*_*j*_ should not be different if two nodes *v*_*i*_ and *v*_*j*_ are adjacent), (b) loss function (predicted scores *f*_*i*_ should be close to the given label of *y*_*i*_). We can obtain predicted score ***f*** by minimizing the following quadratic function:

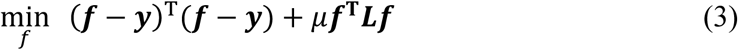

where ***L*** is the graph Laplacian defined as ***L*** = ***D*** − ***W, D*** = diag(*d*_*i*_) is diagonal degree matrix, *d*_*i*_ =∑_*j*_ *w*_*ij*_, and *μ* is user-specific parameter that provides a trade-off between the loss function (first term of Eq. (3)) and smoothness function (second term of Eq. (3)). The closed form of solution ***f*** becomes

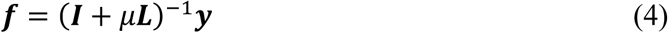

The predicted scores *f* on Eq. (4) can be re-expressed in a block-wise representation by using Eq. (1).^12^

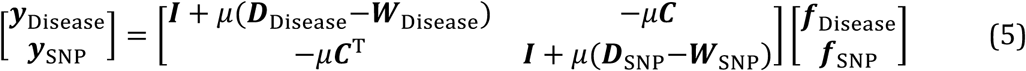

Since the nodes in the SNP-layer are all labeled and nodes in the disease-layer are all unlabeled, Eq. (5) is simplified by substituting ***f***_SNP_ as ***y***_Disease_ and ***y***_Disease_ as ***0***. The predicted scores on the disease-layer are thus obtained as

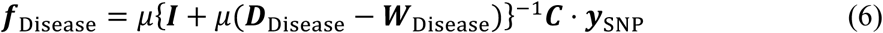

This process is iteratively repeated for each individual patient, and 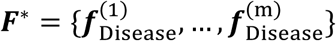 represents the *m*-dimensional comorbidity score vector. To aggregate these vectors, we employ logistic regression as a meta-classifier with **z** ∼ **β**^T^***f***_Disease_ + ***ϵ***. We can then obtain the combined possible comorbidity risk scores as 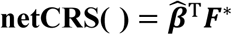 for the individual. A step-by-step process for scoring is summarized with pseudo-code in Supplementary Figure 1.

## 3. Results

In this study, we selected myocardial infarction (PheCode: 411.2) as the index disease of interest. It is commonly known as a heart attack and occurs when blood flow reduces or stops to a part of the heart. Myocardial infarction (MI) is the main undesirable outcome of coronary artery disease. Coronary artery disease, often caused by coronary atherosclerosis, is a common chronic condition characterized by a substantial and complex polygenic contribution to disease risk, with a heritability between 40% and 60%. We describe a MI-specific DS-Net and present comorbidity scores of MI for the individual, **netCRS (***myocardial infarction, MI*).

### 3.1. Experimental Setting

#### 3.1.1. Data for model development and validation set

To build the MI-specific DS-Net and calculate netCRS(MI), a total of 1,403 PheCode-based UK biobank PheWAS summary statistics were obtained from https://www.leelabsg.org/resources.^16^ To construct the myocardial infarction-specific DS-Net, 135 diseases were selected with the following criteria: (a) The diseases were included in the disease-layer if phenotypes had a minimum number of cases larger than 1000, and (b) the diseases were included if phenotypes had at least one shared SNP with myocardial infarction (directly connected with MI). The selected disease categories and disease-layers are described in Figure 2. In the SNP-layer, 39,365 SNPs were selected with genome-wide significance *p*-value threshold ≤ 1 × 10^−4^. Linkage disequilibrium (LD) pruning was performed with thresholds (window size: 50, step size: 5, and *r*^2^ threshold: 0.5). A list of components in the DS-Net is described in Supplementary Table 1.

**Figure 2.**
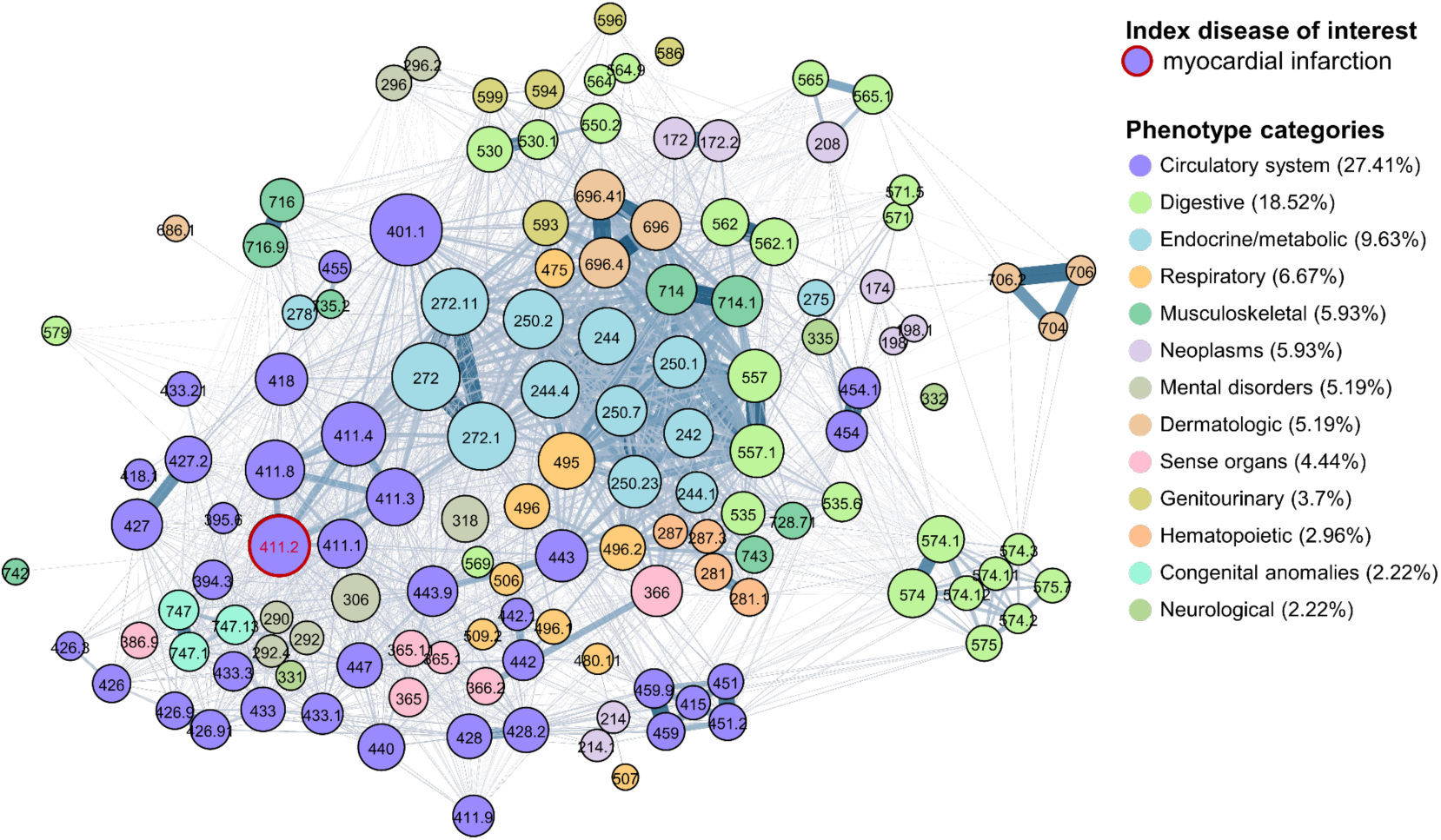
Visualization of MI-specific disease-layer: The node size is the sum of the weighted degree of the node, indicating the relative size, and the node labels represents their PheCode. The thickness of the line represent the edge weights (similarity). Parentheses in disease categories represent the percentages of diseases that belong to a category.

Individual genotype data were collected from the PMBB. The PMBB is an institutional research program that recruits patient-participants throughout the University of Pennsylvania Health System by enrolling at the time of outpatient visits ore more recently, through electronic consenting. Approximately 45,000 of these participants already have genotype data available along with electronic health records (EHR). ICD-9 and ICD-10 codes were aggregated to PheCodes by referring to the PheCode Map 1.2 version.^17-19^ 4,972 individuals of European ancestry were included for this study, all of whom underwent genotyping and had available electronic health record data (Table 1). The detailed genotype QC we performed refers to the previous study ^20^. According to the accumulated medical history at the time of participation, individuals were considered cases for MI if they had at least 2 instances of the PheCode on unique dates, controls if they had no instance of the PheCode, and ‘other/missing’ if they had one instance or a related PheCode. Table 1 describes the list of data and sources for model development and validation cohort.

**Table 1.** Demographics table of the development and validation cohort.

| Development Cohort<br>(Network construction) | UK BioBank PheWAS summary data (UKBB) |  |  |  |  |
| --- | --- | --- | --- | --- | --- |
|  | Phenotypes | 135 (out of 1,403) |  |  |  |
|  | SNPs | 39,365 (after genetic pre-processing) |  |  |  |
| Validation Cohort<br>(Genotype data) | Penn Medicine BioBank (PMBB) |  |  |  |  |
|  |  | Total | MI cases | Controls | p-value |
|  | No. of samples | (N = 4972) | (N = 763) | (N = 4209) |  |
|  | Sex |  |  |  | <0.001 |
|  | Female (%) | 1,854 (37.3%) | 171 (22.4%) | 1683 (40.0%) |  |
|  | Male (%) | 3,118 (62.7%) | 592 (77.6%) | 2526 (60.0%) |  |
|  | Age at enrollment | 62.0 ± 14.8 | 68.4 ± 11.2 | 60.9 ± 15.1 | <0.001 |

#### 3.1.2. Experimental Setting

To evaluate the prediction performance of netCRS using PMBB genotype data, we compared proposed method to PRS with pruning and thresholding (PRS-PT), calculated using PRSice-2^21^. Area under the receiver operating characteristic curve (AUC) was used as performance measure. The model parameters were searched over the following ranges for the respective models. In netCRS(MI), we performed a hyper-parameter search of *μ* for Eq. (4) of graph-based SSL over *μ* = {0.01, 0.1, 1, 10, 100}. The PRS-PT was generated from the sum of the risk alleles weighted by their effect sizes based on GWAS summary statistics from Coronary Artery Disease Genome-wide Replication and Meta-analysis plus the Coronary Artery Disease Genetics (CARDIOGRAMplus C4D consortium).^22^ The parameters were selected from a range of *p*-value thresholds {5 × 10^−8^, 1 × 10^−6^, 0.0001, 0.001, 0.01, 0.05} and LD-based clumping *r*^2^ (0.1 to 0.9) within 1,000 kb. The generated netCRS(MI) and PRS-PT(MI) were compared between MI cases and healthy controls with the logistic regression model, respectively. For both models, the best performance was selected by searching over the respective model-parameter space. The best model of PRS-PT(MI) was determined based on the optimal threshold with the largest Nagelkerke’s *r*^2^ value (in Supplementary Table 1).

#### 3.1.3. Risk predictions of myocardial infarction with netCRS

Table 2 shows the performance comparison of the best PRS-PT(MI) and netCRS(MI) in terms of overall AUC for MI cases and healthy controls. In the results, we included the prediction performance of singleton risk model (netCRS and PRS-PT) and models with covariates of sex and age. We also included the additive models of (PRS-PT + netCRS) with and without covariates. The netCRS with *μ* = {0.1} achieved best predictive performance across both singleton and additive models. When netCRS was used along with the conventional PRS model, the combined model [6] (netCRS + PRS-PT + covariates) achieved an AUC improvement of 28.29%(= (0.7417 − 0.5827)/0.5827) compared to the PRS-PT alone model [1] in MI case prediction. Also, the combined model [6] improved the performance up to 0.7417 of AUC (lifted from 0.6979), comparing to the individual PRS-PT model [4] (AUC improvement of 6.26%). Models with superscript of asterisk were used in further association analysis to validate netCRS and its effectiveness (model [2], [5], and [6])

**Table 2.** Performance comparison of netCRS and PRS-PT in terms of AUC

| Models | Hyper-parameter ($\mu$) for netCRS | | | | |
| --- | --- | --- | --- | --- | --- |
|  | 0.01 | 0.1 | 1 | 10 | 100 |
| [1] PRS-PT | 0.5827 (Baseline) |  |  |  |  |
| [2] netCRS* | 0.6028 | **0.6444** | 0.6395 | 0.6197 | 0.6039 |
| [3] netCRS + PRS-PT | 0.6274 | **0.6609** | 0.6570 | 0.6389 | 0.6255 |
| [4] PRS-PT + Sex + Age | 0.6979 (Baseline) |  |  |  |  |
| [5] netCRS + Sex + Age* | 0.7083 | **0.7287** | 0.7261 | 0.7144 | 0.7051 |
| [6] netCRS + PRS-PT + Sex + Age* | 0.7230 | **0.7417** | 0.7396 | 0.7287 | 0.7199 |

#### 3.1.4. Association analysis of netCRS and PRS

To investigate the effectiveness of the association between both risk scoring models and covariates with age and sex, we assessed multiplicative interactions between netCRS and each of the stratification variables. We stratified participants based on quartiles of netCRS; low risk (0th-25th), intermediate risk (26th-50th), high risk (51st-75th), and very high risk (76th-100th). Compared with the low-netCRS risk group, the higher netCRS risk group had higher odds ratios in the validation cohort. In stepwise multivariate models (model [5] and [6]), the models with covariates and/or PRS-PT remained significantly (Table 3). Participants in the very high-netCRS risk group for MI had approximately four-fold increased risk of MI occurrence relative to those with the corresponding low-genetic risk group (shown in Table 3). In addition, we investigated the benefit of using netCRS and PRS together in screening high-risk groups for MI. Table 4 demonstrates that combinations of MI-PRS and netCRS were able to capture the risk of MI up to approximately eight-fold higher than the low-risk group. Supplementary Table 3 provides demographics of participants according to netCRS risk groups.

**Table 3.**
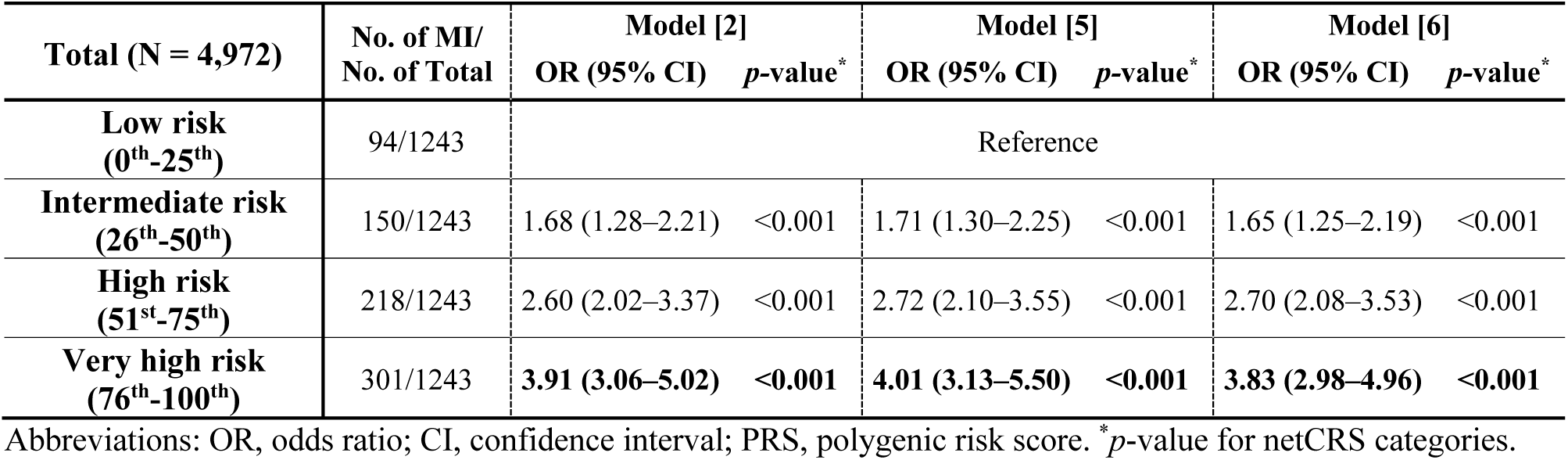
Diagnostic odds ratio and 95% confidential intervals for the MI according to netCRS risk group: We compared three different models: (a) model [2]: netCRS alone, (b) model [5]: netCRS + sex + age, and (c) model [6]: netCRS + PRS-PT + sex + age.

| Total (N = 4,972) | No. of MI/<br>No. of Total | Model [2] |  | Model [5] |  | Model [6] |  |
| --- | --- | --- | --- | --- | --- | --- | --- |
|  |  | OR (95% CI) | *p*-value* | OR (95% CI) | *p*-value* | OR (95% CI) | *p*-value* |
| Low risk<br>(0<sup>th</sup>-25<sup>th</sup>) | 94/1243 | Reference |  |  |  |  |  |
| Intermediate risk<br>(26<sup>th</sup>-50<sup>th</sup>) | 150/1243 | 1.68 (1.28–2.21) | <0.001 | 1.71 (1.30–2.25) | <0.001 | 1.65 (1.25–2.19) | <0.001 |
| High risk<br>(51<sup>st</sup>-75<sup>th</sup>) | 218/1243 | 2.60 (2.02–3.37) | <0.001 | 2.72 (2.10–3.55) | <0.001 | 2.70 (2.08–3.53) | <0.001 |
| Very high risk<br>(76<sup>th</sup>-100<sup>th</sup>) | 301/1243 | **3.91 (3.06–5.02)** | **<0.001** | **4.01 (3.13–5.50)** | **<0.001** | **3.83 (2.98–4.96)** | **<0.001** |

Abbreviations: OR, odds ratio; CI, confidence interval; PRS, polygenic risk score. \**p*-value for netCRS categories.

**Table 4.** Genetic subgroups based on the combinations of PRS and netCRS

| Odds ratio*<br>(No. of MI / No. of Total) |  | PRS-PT(MI) |  |  |  |
| --- | --- | --- | --- | --- | --- |
|  |  | Low risk<br>(0<sup>th</sup>-25<sup>th</sup>) | Intermediate risk<br>(26<sup>th</sup>-50<sup>th</sup>) | High risk<br>(51<sup>st</sup>-75<sup>th</sup>) | Very high risk<br>(76<sup>th</sup>-100<sup>th</sup>) |
| netCRS(MI) | Low risk (0<sup>th</sup>-25<sup>th</sup>) | Reference<br>(19/334) | 1.18<br>(20/299) | 1.35<br>(21/273) | 2.46<br>(34/243) |
|  | Intermediate risk (26<sup>th</sup>-50<sup>th</sup>) | 1.46<br>(23/276) | 2.36<br>(36/268) | 2.77<br>(45/286) | 3.07<br>(46/263) |
|  | High risk (51<sup>st</sup>-75<sup>th</sup>) | 2.07<br>(33/280) | 4.59<br>(71/272) | 3.94<br>(52/241) | 4.55<br>(60/232) |
|  | Very high risk (76<sup>th</sup>-100<sup>th</sup>) | 4.04<br>(52/226) | 4.66<br>(58/219) | 5.60<br>(78/245) | **7.88**<br>**(113/252)** |

\*For calculating odds ratio, we performed multivariate logistic regression analysis for MI classification task (myocardial infarction (MI) cases versus Normal control). Logistic model: (MI cases vs. Normal control) ~ 16 combinations (PRS and netCRS groups) + sex + age. With the lowest risk group (Low PRS group & Low netCRS group) as a reference, the odds ratio of each combination was reported in this table.

## 4. Conclusion

In this study, we developed and proposed netCRS, a network-based disease comorbidity risk scoring algorithm based upon biobank-scale PheWAS summary statistics. To improve the prediction ability of PRS, we introduced a novel combined comorbidity risk scores using a multi-layered network. Most current biological networks suggest only associative information between biological components according to aggregated population-level data ^23^. Although these population-level networks provide insights regarding the interaction of components, it is not easy to obtain individual inference from them.

To solve this problem, we proposed a novel method for the prediction of individual-level risk scores from population-level interactome. We first constructed a DDN (disease-layer) which elaborates on the genetic associations among multiple phenotypes in UKBB PheWAS data. In order to use the disease-layer at the individual-level, we attached a SNP-layer to the disease-layer. The final developed network is a disease-SNP heterogeneous multi-layered network denoted as DS-Net. We employed graph-based SSL on the network to devise a network-based scoring algorithm. The SNP-layer is a single network that serves as initial labeling to receive individual genotyping data, and the disease-layer is an output network. The disease-layer serves as the predicted possible comorbidity risk scores in which the individual’s genotype is propagated. To obtain layer-wise predicted scores, a layer-wise positive-unlabeled learning setting was employed, where the all nodes on the disease-layer are unlabeled and all the SNPs on the SNP-layer are labeled. Graph-based SSL can operate in this problem setting to propagate label information according to the topology of the network. The resulting netCRS is an estimated comorbidity score that integrates pre-defined genetic association between phenotypes using the underlying structure of the DS-Net. This score includes not only genetic information about a specific target disease, but also multiple associations of diseases. We validated the proposed netCRS by considering MI as index disease of interest. The netCRS model outperformed the conventional PRS-PT model in predicting MI patients and healthy controls. From experimental results of the association analysis, it is noteworthy that netCRS and PRS-PT work complementary to one another in identifying the very high-risk group of patients with myocardial infarction.

The current proposed method still has room for improvement. First, when constructing a disease-specific heterogeneous multi-layered network, it is expected that better comorbidity scores will be obtained if more precise criteria are applied to node selection. Second, our network was constructed using only common variants from PheWAS summary data. If we expand the network to include rare variants and other clinical information, we expect that using these risk prediction models will allow for the development of prevention strategies and reduction of MI morbidity and mortality. Also, the current disease-layer was constructed according to shared common SNPs between diseases. We can also try to build the DDN using different forms of genetic correlations such as LD regression scores. For future work, we will test netCRS in various diseases and compare netCRS with more recent PRS approaches in order to prove its generalized prediction performance.

## 5. Acknowledgments

This work was supported by NIGMS R01 GM138597, NLM R01 NL012535, and S10OD023495. We thank the staff of the Regeneron Genetics Center for DNA sequencing from PMBB participants. Use of the UK Biobank Resource in the current study was approved under Application Number 67855. Supplementary data are available at https://github.com/dokyoonkimlab/netCRS/blob/main/netCRS_supplemental.pdf

## Supplementary documents

**Supplementary Figure 1.**
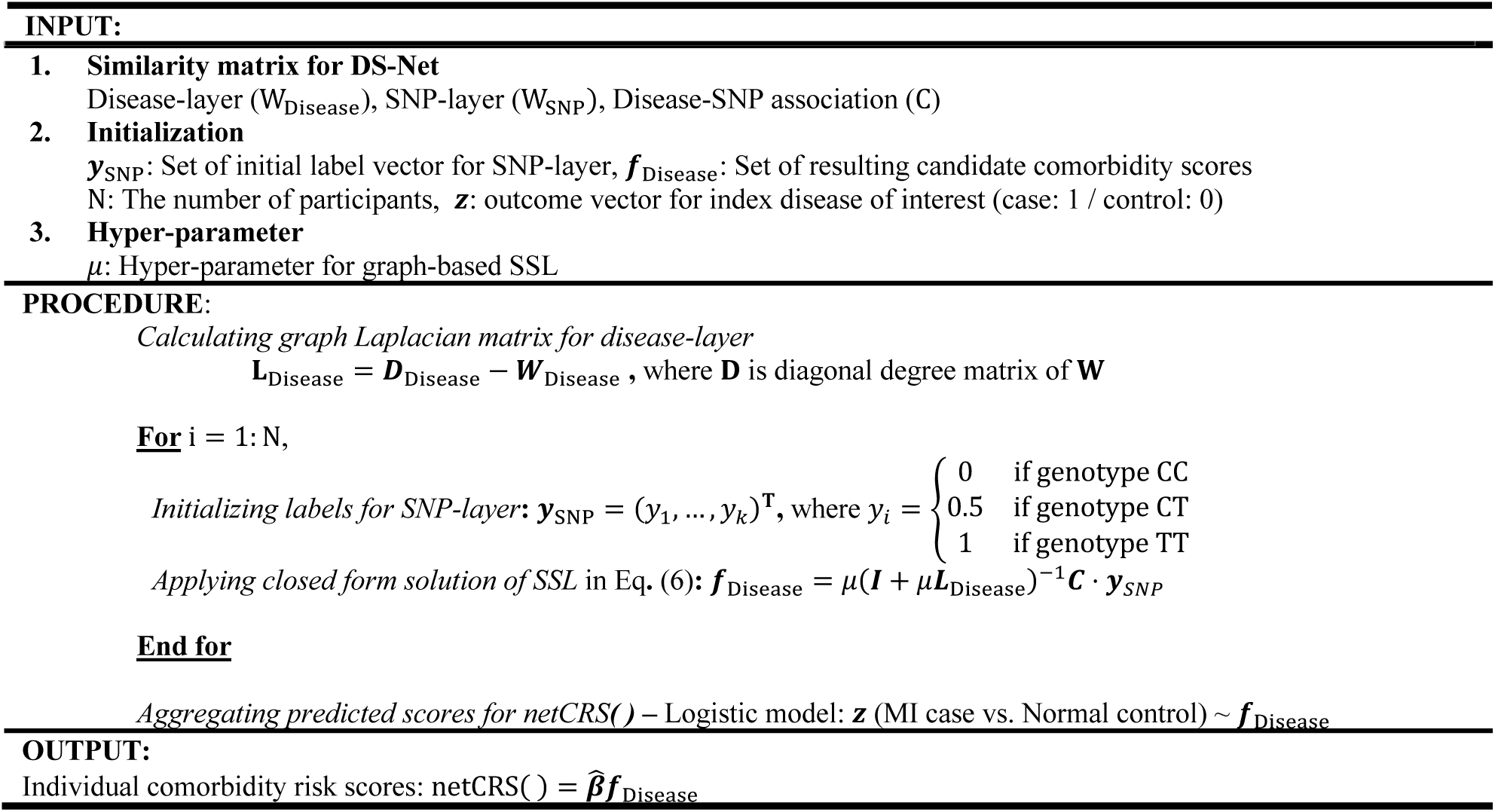
Pseudo-code for netCRS of section 2.2

**Supplementary Table 1.**
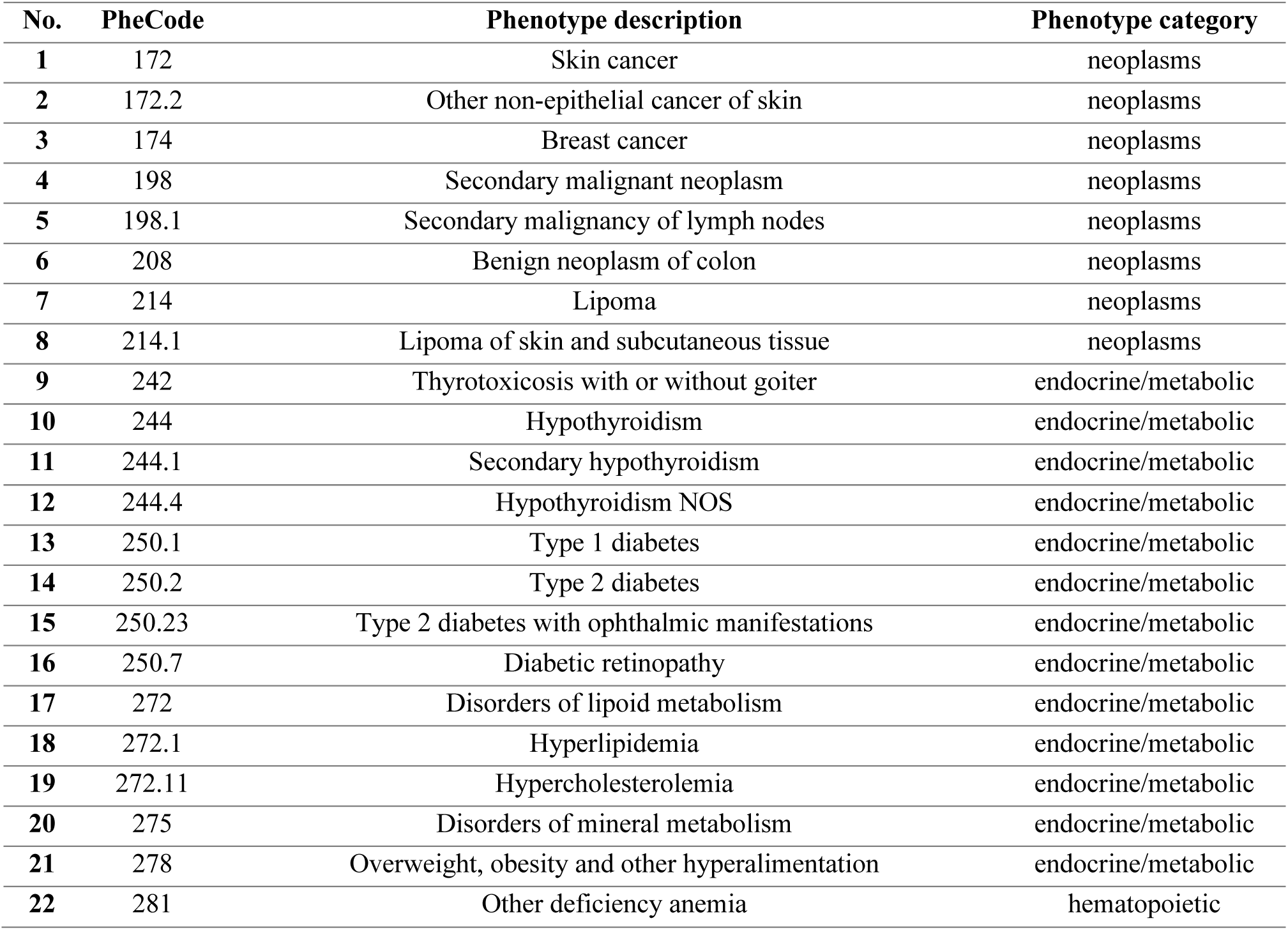
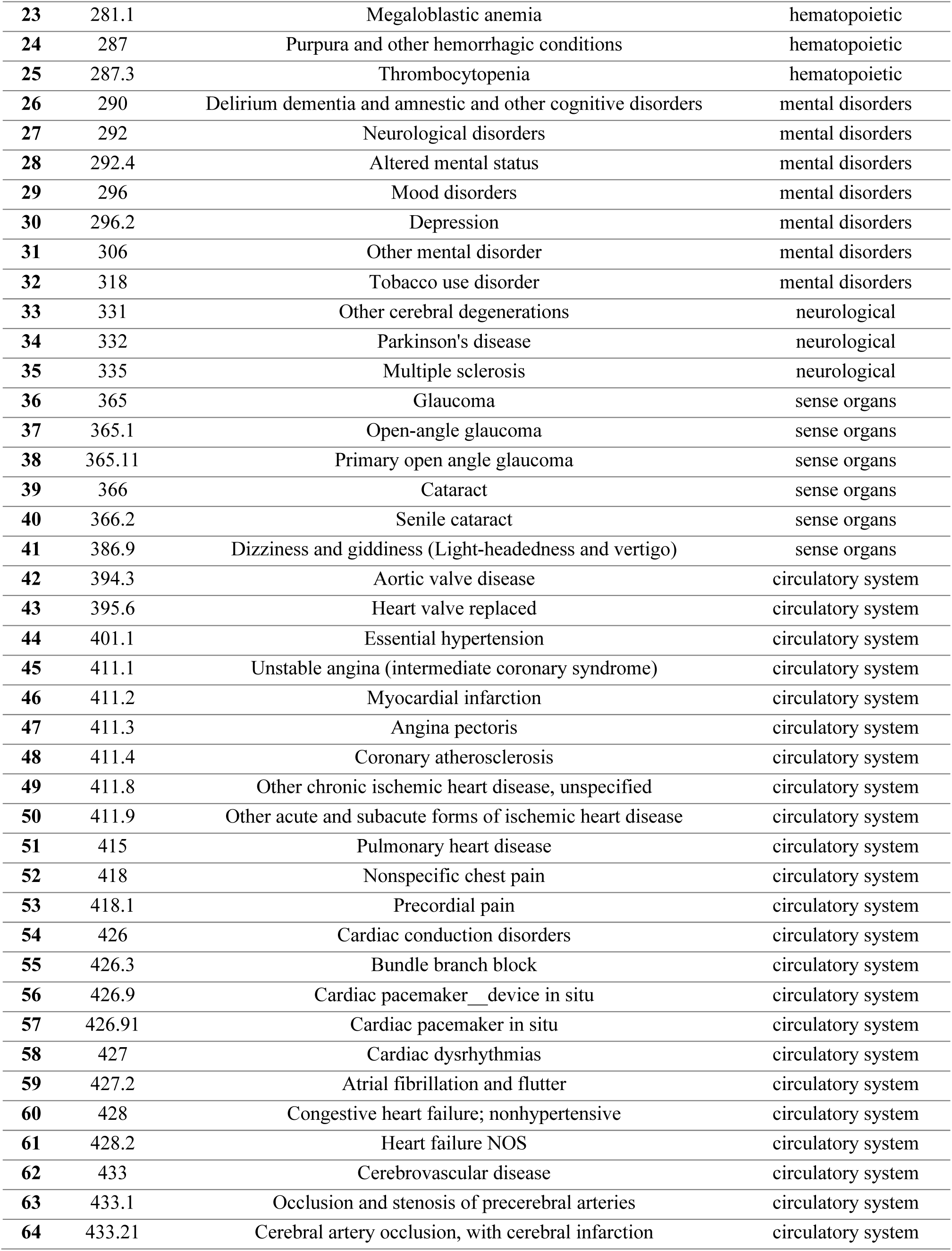
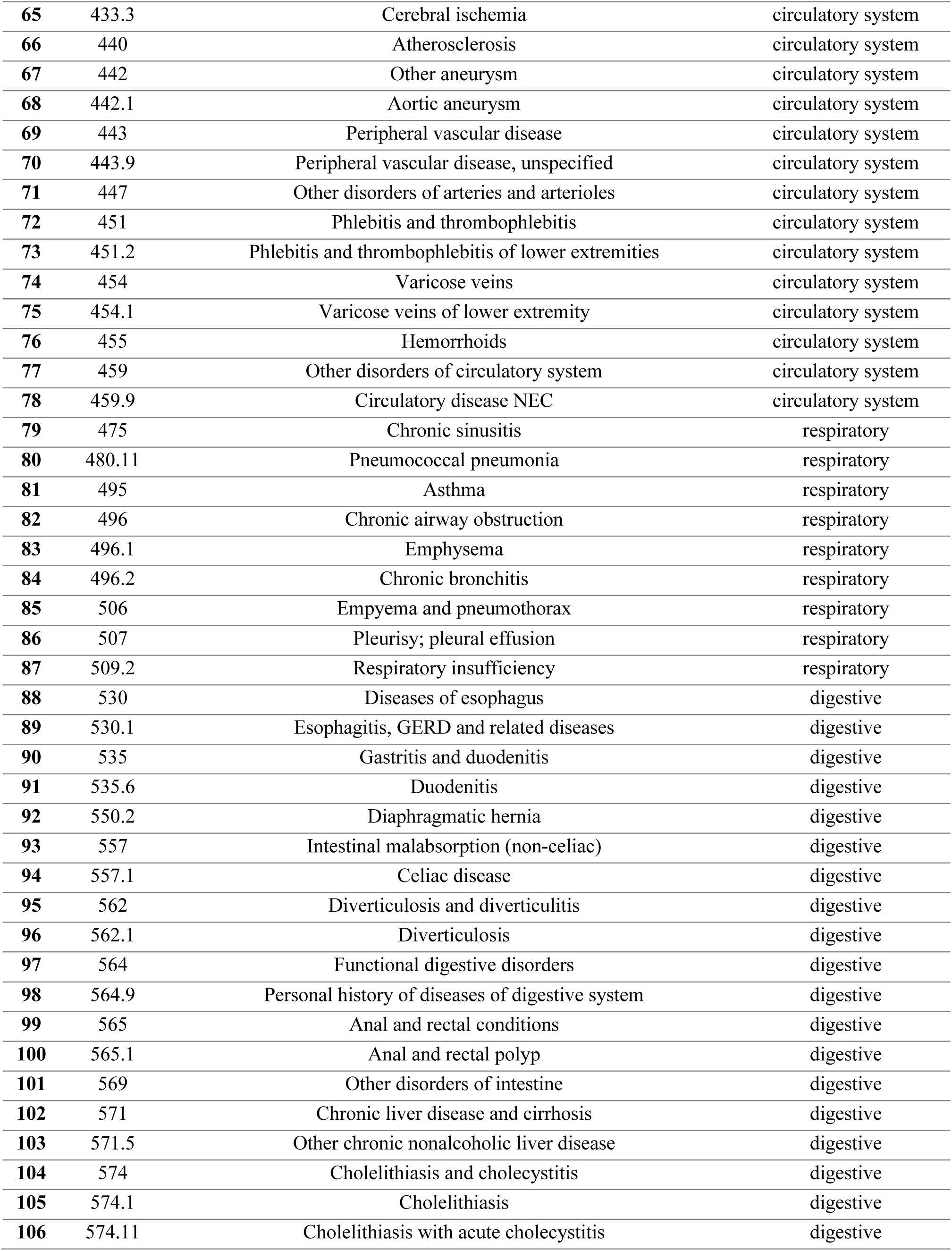
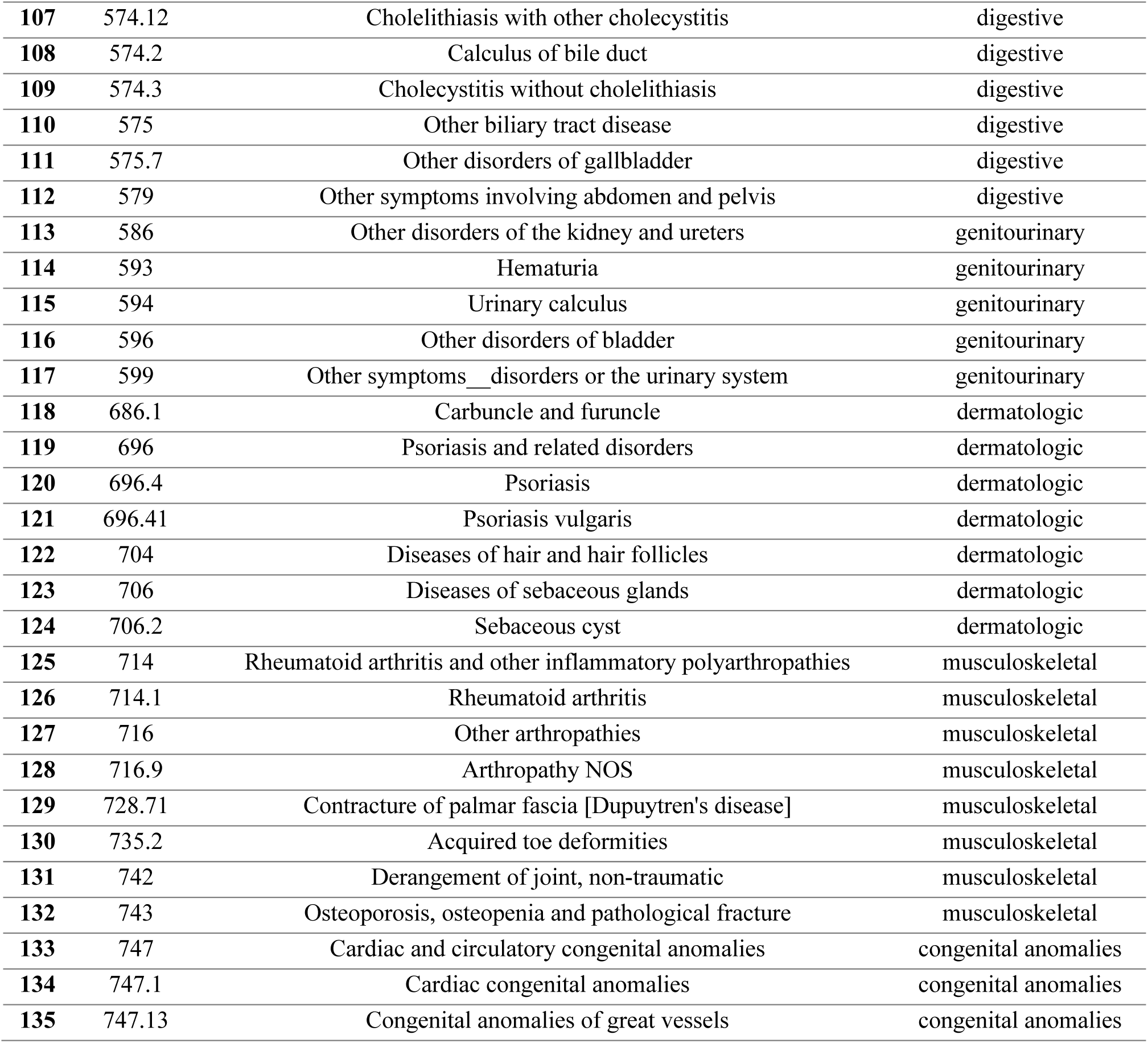
A list of phenotypes in disease-layer.

**Supplementary Table 2.** Nagelkerke’s *r*^2^ values of PRS in the classification of myocardial infarction cases and controls at the linkage disequilibrium clumping and *p*-value thresholds

| LD-clump $r^2$ threshold | $p$-value threshold | Nagelkerke's $R^2$ | $p$-value | Beta | Standard error | Number of used SNPs |
| --- | --- | --- | --- | --- | --- | --- |
| >0.1 | 5.00E-08 | 0.0093 | 8.00E-14 | 0.5414 | 0.0725 | 32 |
|  | 1.00E-06 | 0.0085 | 8.53E-13 | 0.4108 | 0.0574 | 63 |
|  | 0.0001 | 0.0111 | 3.52E-16 | 0.2882 | 0.0353 | 307 |
|  | 0.001 | 0.0111 | 4.30E-16 | 0.1609 | 0.0198 | 1483 |
|  | 0.01 | 0.0099 | 1.52E-14 | 0.0708 | 0.0092 | 9685 |
|  | 0.05 | 0.0085 | 9.78E-13 | 0.0366 | 0.0051 | 37482 |
| >0.2 | 5.00E-08 | 0.0071 | 6.30E-11 | 0.3831 | 0.0586 | 39 |
|  | 1.00E-06 | 0.0070 | 9.55E-11 | 0.3143 | 0.0485 | 73 |
|  | 0.0001 | 0.0095 | 5.07E-14 | 0.2287 | 0.0304 | 348 |
|  | 0.001 | 0.0110 | 5.66E-16 | 0.1418 | 0.0175 | 1638 |
|  | 0.01 | 0.0116 | 9.44E-17 | 0.0678 | 0.0082 | 10863 |
|  | 0.05 | 0.0103 | 4.74E-15 | 0.0348 | 0.0044 | 44067 |
| >0.3 | 5.00E-08 | 0.0070 | 9.82E-11 | 0.3126 | 0.0483 | 45 |
|  | 1.00E-06 | 0.0075 | 2.31E-11 | 0.2684 | 0.0401 | 85 |
|  | 0.0001 | 0.0103 | 4.42E-15 | 0.2024 | 0.0258 | 388 |
|  | 0.001 | 0.0117 | 7.15E-17 | 0.1274 | 0.0153 | 1805 |
|  | 0.01 | 0.0130 | 1.58E-18 | 0.0640 | 0.0073 | 11911 |
|  | 0.05 | 0.0111 | 4.27E-16 | 0.0316 | 0.0039 | 49675 |
| >0.4 | 5.00E-08 | 0.0063 | 7.37E-10 | 0.2552 | 0.0414 | 50 |
|  | 1.00E-06 | 0.0065 | 4.72E-10 | 0.2072 | 0.0333 | 99 |
|  | 0.0001 | 0.0093 | 1.03E-13 | 0.1647 | 0.0221 | 438 |
|  | 0.001 | 0.0115 | 1.15E-16 | 0.1123 | 0.0136 | 1962 |
|  | 0.01 | 0.0144 | 2.51E-20 | 0.0603 | 0.0065 | 12985 |
|  | 0.05 | 0.0121 | 2.38E-17 | 0.0292 | 0.0034 | 55003 |
| >0.5 | 5.00E-08 | 0.0061 | 1.84E-09 | 0.2139 | 0.0356 | 56 |
|  | 1.00E-06 | 0.0063 | 7.36E-10 | 0.1746 | 0.0283 | 113 |
|  | 0.0001 | 0.0088 | 4.52E-13 | 0.1361 | 0.0188 | 492 |
|  | 0.001 | 0.0118 | 5.29E-17 | 0.1006 | 0.0120 | 2133 |
|  | 0.01 | 0.0153 | 1.42E-21 | 0.0557 | 0.0058 | 14160 |
|  | 0.05 | 0.0128 | 2.60E-18 | 0.0267 | 0.0031 | 60663 |
| >0.6 | 5.00E-08 | 0.0057 | 6.01E-09 | 0.1700 | 0.0292 | 67 |
|  | 1.00E-06 | 0.0064 | 6.34E-10 | 0.1549 | 0.0251 | 124 |
|  | 0.0001 | 0.0092 | 1.23E-13 | 0.1202 | 0.0162 | 561 |
|  | 0.001 | 0.0121 | 1.99E-17 | 0.0888 | 0.0105 | 2353 |
|  | 0.01 | 0.0162 | 1.05E-22 | 0.0506 | 0.0052 | 15489 |
|  | 0.05 | 0.0134 | 4.72E-19 | 0.0240 | 0.0027 | 66890 |
| >0.7 | 5.00E-08 | 0.0059 | 3.34E-09 | 0.1499 | 0.0253 | 76 |
|  | 1.00E-06 | 0.0062 | 1.30E-09 | 0.1265 | 0.0208 | 146 |
|  | 0.0001 | 0.0089 | 3.51E-13 | 0.0990 | 0.0136 | 631 |
|  | 0.001 | 0.0118 | 5.97E-17 | 0.0733 | 0.0088 | 2650 |
|  | 0.01 | 0.0170 | 9.14E-24 | 0.0447 | 0.0044 | 17122 |
|  | 0.05 | 0.0148 | 6.25E-21 | 0.0219 | 0.0023 | 74227 |
| >0.8 | 5.00E-08 | 0.0049 | 7.42E-08 | 0.0839 | 0.0156 | 100 |
|  | 1.00E-06 | 0.0055 | 8.75E-09 | 0.0791 | 0.0138 | 188 |
|  | 0.0001 | 0.0084 | 1.37E-12 | 0.0702 | 0.0099 | 757 |
|  | 0.001 | 0.0124 | 8.29E-18 | 0.0592 | 0.0069 | 3042 |
|  | 0.01 | 0.0170 | 8.31E-24 | 0.0377 | 0.0057 | 19321 |
|  | 0.05 | 0.0160 | 2.19E-22 | 0.0191 | 0.0020 | 84126 |
| >0.9 | 5.00E-08 | 0.0036 | 3.72E-06 | 0.0454 | 0.0098 | 135 |
|  | 1.00E-06 | 0.0046 | 1.50E-07 | 0.0472 | 0.0090 | 247 |
|  | 0.0001 | 0.0071 | 6.79E-11 | 0.0448 | 0.0069 | 969 |
|  | 0.001 | 0.0112 | 3.59E-16 | 0.0406 | 0.0050 | 3747 |
|  | **0.01** | **0.0171** | **7.40E-24** | **0.0282** | **0.0028** | **23106** |
|  | 0.05 | 0.0166 | 3.65E-23 | 0.0152 | 0.0015 | 100493 |

**Supplementary Table 3.** Demographics of participants according to netCRS risk groups

| netCRS |  | Low | Intermediate | High | Very high | *P*-value |
| --- | --- | --- | --- | --- | --- | --- |
| Number of samples |  | (n=1243) | (n=1243) | (n=1243) | (n=1243) |  |
| Age |  | 61.7 ± 15.2 | 61.8 ± 15.2 | 62.0 ± 14.7 | 62.6 ± 14.3 | 0.116 |
| Sex | Female (%) | 778 (62.6%) | 771 (62.0%) | 769 (61.9%) | 800 (64.4%) | 0.539 |
| Myocardial infarction | Cases (%) | 94 (7.6%) | 150 (12.1%) | 218 (17.5%) | 301 (24.20%) | <0.001 |
|  | Controls (%) | 1149 (92.4%) | 1093 (87.9%) | 1025 (82.5%) | 942 (75.80%) |  |

